# Unexpected similarity between HIV-1 reverse transcriptase and tumor necrosis factor revealed by binding site image processing

**DOI:** 10.1101/2021.06.09.447723

**Authors:** Merveille Eguida, Didier Rognan

## Abstract

Rationalizing the identification of hidden similarities across the repertoire of druggable protein cavities remains a major hurdle to a true proteome-wide structure-based discovery of novel drug candidates. We recently described a new computational approach (ProCare), inspired by numerical image processing, to identify local similarities in fragment-based subpockets. During the validation of the method, we unexpectedly identified a possible similarity in the binding pockets of two unrelated targets, human tumor necrosis factor alpha (TNF-α) and HIV-1 reverse transcriptase (HIV-1 RT). Microscale thermophoresis experiments confirmed the ProCare prediction as two of the three tested and FDA-approved HIV-1 RT inhibitors indeed bind to soluble human TNF-α trimer. Interestingly, the herein disclosed similarity could be revealed neither by state-of-the-art binding sites comparison methods nor by ligand-based pairwise similarity searches, suggesting that the point cloud registration approach implemented in ProCare, is uniquely suited to identify local and unobvious similarities among totally unrelated targets.

**AUTHOR SUMMARY:** Computational comparison of binding sites in proteins can provide insights on potential unrelated proteins that may bind to similar ligands. However, accurate prediction of binding site similarity requires powerful methods, ideally able to detect even local similarities. We herewith applied a recently developed computer vision method to identify an unexpected binding site similarity between two totally unrelated proteins that was confirmed experimentally by in vitro biophysical binding assays. Considering more precisely local similarities can therefore efficiently guide drug discovery, notably to repurpose existing drug candidates.

## INTRODUCTION

Among the many possible approaches to structure-based drug design [1,2], inferring novel ligand properties from the large-scale comparison of their possible binding pockets gains popularity as the repertoire of protein cavities of known three-dimensional (3D) structures (pocketome) is constantly increasing, thereby offering unique opportunities to design ligands while simultaneously considering multiple targets [3]. The term ‘pocketome’ was first coined in 2004 by An et al. [4] to describe the universe of cavities located at the surface of macromolecules and capable of binding low molecular-weight ligands. A systematic survey of currently available three-dimensional structures [5], suggests that its size is estimated to ca. 250,000 pockets [6] out of which 10-15% are accommodating true drug-like compounds [7,8]. Pocket locations can be inferred from the position of already-bound molecules or predicted on the fly, by one of the many available cavity detection methods [3,9]. The pockeome space can then be searched by numerous computational tools [10] for similarity to any query cavity to predict evolutionary relationships and protein-ligand interactions [3]. The later application is notably of paramount importance to the drug discovery field as it may reveal hidden relationships for guiding the design of safer drug candidates with a precise control of selectivity [3] with respect to either on-targets (polypharmacology approach) [11] or off-targets (side effects mitigation) [12], in a time and cost-effective manner [13].

Currently available methods are generally able to detect global similarities between two druggable pockets from different proteins, and therefore permit to transfer drug-like compounds from one target space to another [3]. Identifying more subtle local similarities at the level of fragment-bound pockets remains a much more difficult problem [14] as it requires a searchable archive of fragment-bound subpockets [15–17] and a computational method focusing on local subpocket descriptors. Consequently, there are still very few reports of experimentally verified subpocket similarity examples that have enabled the transfer of chemical fragments across unrelated proteins [18]. To fill the need for local similarity searching methods while comparing pockets of different sizes, we developed a novel method (ProCare) [17] relying on point cloud registration, a numerical image processing to find a spatial transformation (*e.g*., scaling, rotation and translation) that aligns two point clouds [19, 20]. ProCare uses as input a point cloud representation of the protein pocket or subpockets, where each point is annotated by eight possible pharmacophoric features complementary to that of the pocket microenvironment [21]. Since ProCare uses local descriptors to compare and align binding subpockets, the method is particularly suited to fragment-based design strategies aimed at positioning fragments in subpockets of any druggable cavity.

While validating the method by focused benchmarking studies [17], we noticed some unexpected local similarity between subpockets from two unrelated proteins: the asymmetric human tumor necrosis factor alpha (TNF-α) trimer [22] and the human immunodeficiency virus type 1 reverse transcriptase (HIV-1 RT) [23]. To exclude potential artifacts or biases and provide a strong statistical support to this initial prediction, we here systematically compared the inner cavity of three inhibitor-bound TNF-α trimer structures with 122 non-nucleoside inhibitor-bound HIV-1 RT X-ray structures. In a large majority of pairwise comparisons, the corresponding subpockets were deemed similar, a prediction that could be confirmed by biophysical experiments evidencing a direct micromolar binding of non-nucleoside HIV-1 RT inhibitors to human soluble TNF-α. Interestingly, this unexpected similarity could not be recovered by state-of-the-art cavity comparisons tools suggesting the unique ability of ProCare to delineate subtle local relationships between unrelated target cavities.

## RESULTS AND DISCUSSION

Identifying similarity between pockets from different proteins suggests that the latter might bind to similar molecules. Although molecular recognition is a dynamic and complex process, the above hypothesis is worth investigating in drug design for hit discovery or for potential off-targets detection. We previously described ProCare [17], a novel computational method relying on a point cloud registration algorithm [19,20] to assess the similarity between protein pockets. ProCare computes and uses local descriptors, which makes it particularly suitable for detecting local similarities among cavities of different sizes. Typically, ProCare aligns the cavities, described by a cloud of 3D points labeled with pharmacophoric features, by comparing the point descriptors and then derives a similarity score. While benchmarking the ProCare method by comparing protein structures disclosed for the first time to druggable sc-PDB cavities [8], we noticed unexpected high similarities between the core pocket at the interface of the asymmetric human TNF-α trimer [22], and several non-nucleoside binding sites of inhibitor-bound HIV-1 reverse transcriptase (**S1 Table**). To further investigate that hypothesis, exclude a potential bias in the ProCare alignment/scoring method and give a stronger statistical support to our prediction, we systematically compared three binding sites from available asymmetric human TNF-α X-ray structures [22] to that of 122 HIV-1 RT structures bound to non-nucleoside inhibitors.

### Comparison of TNF-α trimer and HIV-1 reverse transcriptase binding sites

A ProCare similarity matrix was built by comparing cavities of three asymmetric TNF-α structures (PDB identifiers 6OOY, 6OOZ and 6OP0) co-crystallized with benzimidazole ihibitors to 122 cavities (yielding 195 subpockets) of non-nucleoside HIV-1 RT inhibitors (**Fig 1**). We observed that 76% of all pairwise comparisons were scored higher than the previously determined ProCare similarity score threshold of 0.47 (**Fig 1A**). To exclude the possibility that the predicted similarity is caused by peculiar mutations of the HIV-1 RT non-nucleoside biding site, we also compared pairwise similarities for both wild type and mutated HIV-1 RT pockets, but did not observe significant differences in the percentage of HIV-1 RT pockets predicted similar to that of TNF-α (74% and 82% of similar pockets for wild type and mutants, respectively). We thus conclude that the predicted similarity between pockets from these two unrelated targets is independent on the chosen PDB structures and is not biased by mutations in the HIV-1 RT binding site.

**Fig 1.**
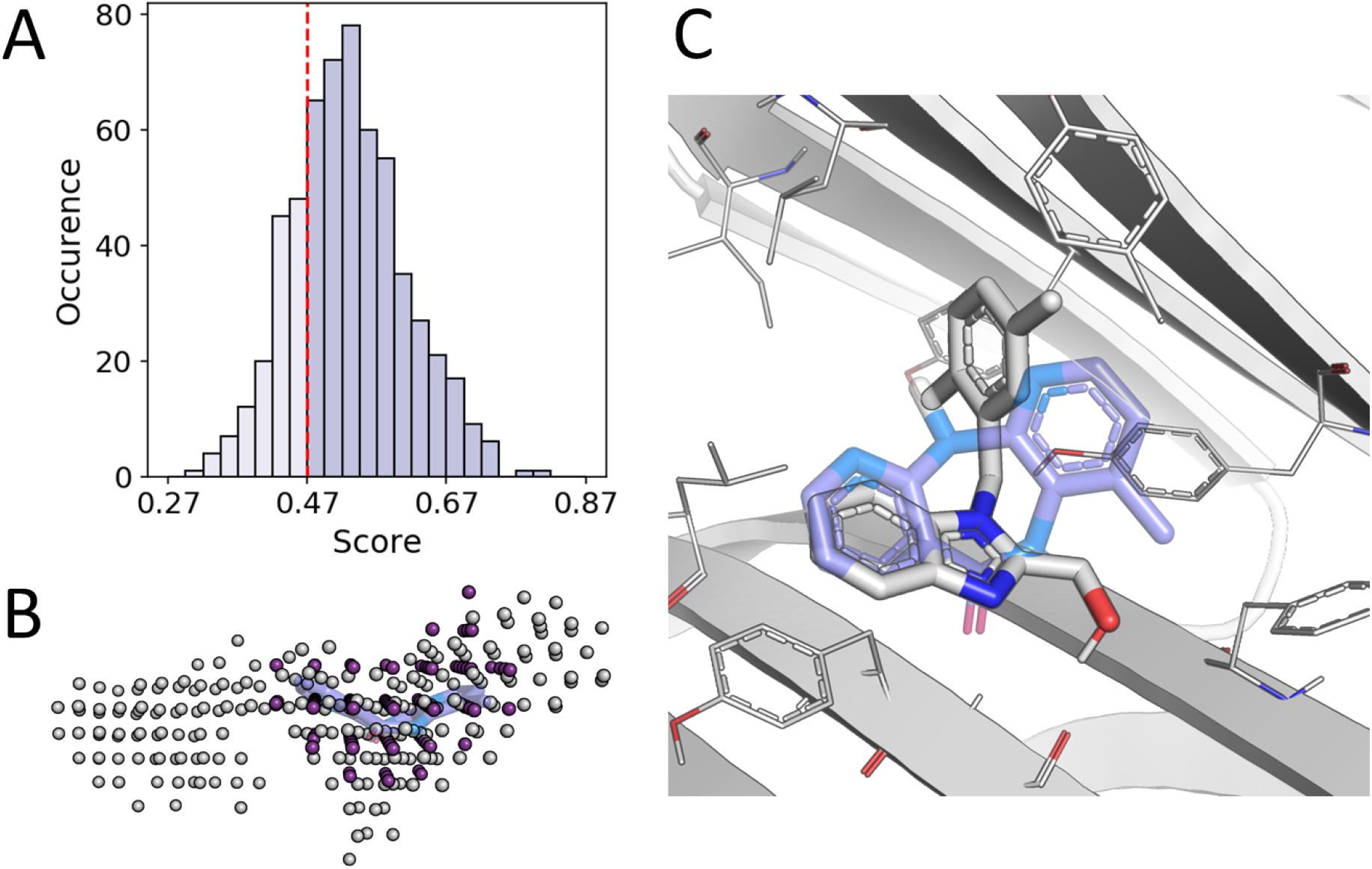
Comparison of TNF-α trimer and HIV-1 RT binding sites with ProCare. (**A**) Distribution of pairwise similarity scores. Entries scoring above 0.47 (threshold marked by the red dashed line) are considered similar. (**B**) Alignment of nevirapine (NVP) subpocket (PDB ID: 1VRT, HET code: NVP, color: purple) to the TNF-α pocket (PDB ID: 6OOY, HET code: A7M, color: white). A video of the alignment is available as supplementary information (**S1 Video**). (**C**) Corresponding alignment of NVP (carbon atoms, purple sticks) in the TNF-α binding site after applying the previous transformation. Carbon atoms of the TNF-α inhibitor are shown as white sticks. Nitrogen and oxygen atoms are colored in blue and red, respectively.

Since ProCare yields 3D alignments of the compared objects (subpockets onto the target pockets), we could visually inspect the rationale behind the alignments. As an example, the alignment of a nevirapine-bound subpocket in HIV-1 RT (PDB ID 1VRT) to one TNF-α cavity (PDB ID 6OOY) captures a local shape similarity in pockets accommodating butterfly-shaped ligands [24] (**Fig 1B**). Applying the ProCare-derived transformation matrix to the HIV-1 RT ligand yielded a coherent shape alignment of the ligands where the two aromatic moieties of the nevirapine tricyclic fragment are matching the two benzene moieties in the TNF-α inhibitor (**Fig 1C**). Accordingly, H-bond donor groups (amide oxygen in nevirapine, benzimidazole nitrogen atom in the TNF-α inhibitor) were also matched. In order to prioritize HIV-1 RT inhibitors for experimental validation of our hypothesis, we checked which inhibitors were bound to the HIV-RT subpockets that are predicted the closest to the TNF-α cavity (**Table 1**).

**Table 1.** Bound inhibitors of the HIV-1 reverse transcriptase cavities found similar to TNF-α cavities

| HIV-RT Inhibitor<sup>a</sup> | HIV1-RT PDB entry | TNF-α PDB entry | ProCare score | Rank |
| --- | --- | --- | --- | --- |
| NNI | 2VG7 | 6OOZ | 0.810 | 1 |
| EFZ (Efavirenz) | 1FKO | 6OOZ | 0.773 | 2 |
| NVP (Nevirapine) | 1LWC | 6OOZ | 0.737 | 3 |
| TNK | 1S1V | 6OOZ | 0.731 | 4 |
| NVP (Nevirapine) | 2HNY | 6OOZ | 0.729 | 5 |
| ... |  | ... | ... | ... |
| SPP (Delavirdine) | 1KLM | 6OOZ | 0.484 | 408 |

<sup>a</sup> PDB chemical component identifier (Drug name in brackets).

Among the corresponding inhibitors, two easily purchasable FDA-approved drugs (efavirenz, neviparine; **Fig 2**) are almost entirely buried in the HIV-1 RT subpockets found similar to the TNF-α cavity, exhibit a size and molecular volume similar to that of two TNF-α inhibitors (A7M, A6Y; **Fig 2**) and were therefore selected for biological evaluation. In addition, we also considered a third marketed inhibitor (delavirdine; **Table 1, Fig 2**) whose pocket was found much less similar to that of TNF-α, although just above the 0.47 ProCare similarity threshold.

**Fig 2.**
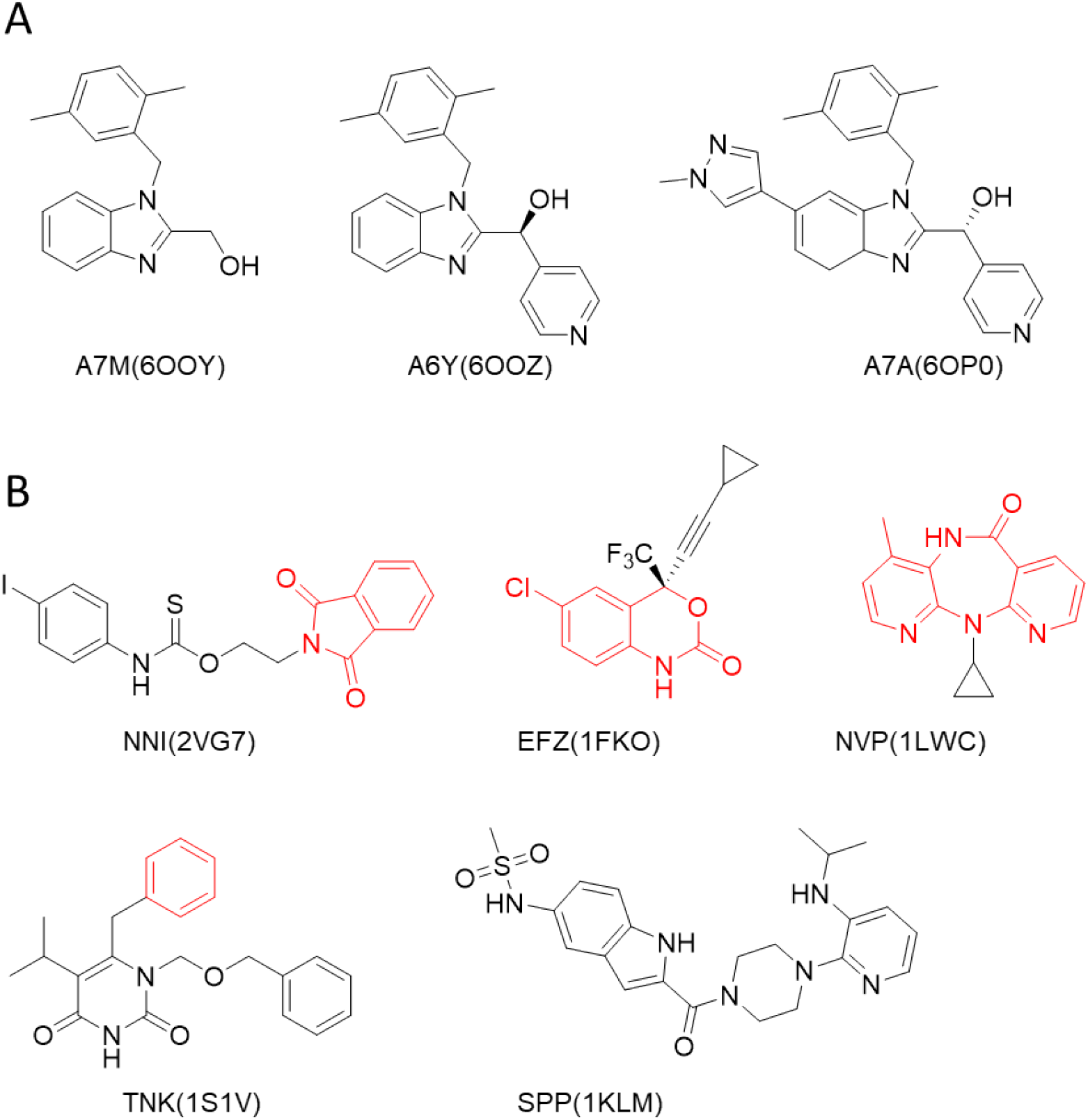
Structures of TNF-α and HIV-1 RT non-nucleoside inhibitors. (**A**) TNF-α inhibitors and (**B**) HIV-1 RT non-nucleoside inhibitors. Inhibitors are named according to their PDB chemical component identifier and PDB entry (between brackets). Red substructures indicate the fragment binding to the HIV-1 RT subpocket found similar to the TNF-α cavity. Please note that delavirdine (PDB HET code SPP) is not fragmented by the IChem fragmentation tool [25].

### Non-nucleoside HIV-1 RT inhibitors bind to human TNF-α

According to the similarity principle, similar ligands may bind to similar binding sites. We then hypothesized that some reverse transcriptase non-nucleoside inhibitors may be able to bind to the central cavity of the TNF-α trimer. Three different non-nucleoside FDA-approved drugs (nevirapine, efavirenz and delavirdine) were thus tested for direct binding to a fluorescent-labelled TNF-α trimer by microscale thermophoresis (MST), a robust and sensitive biophysical method to detect and quantify molecular interactions in solution [26,27]. The MST signal is based on ligand-dependent temperature-induced changes (thermophoresis, temperature-related fluorescence intensity) of the fluorescence emission of the labelled protein target. The 17.3 kDa homotrimeric TNF-α that spontaneously assemble in solution [28,29] was therefore labelled by a RED-fluorescent probe for MST experiments in presence of increasing concentrations of the three HIV-1 RT inhibitors (**Fig 3**).

**Fig 3.**
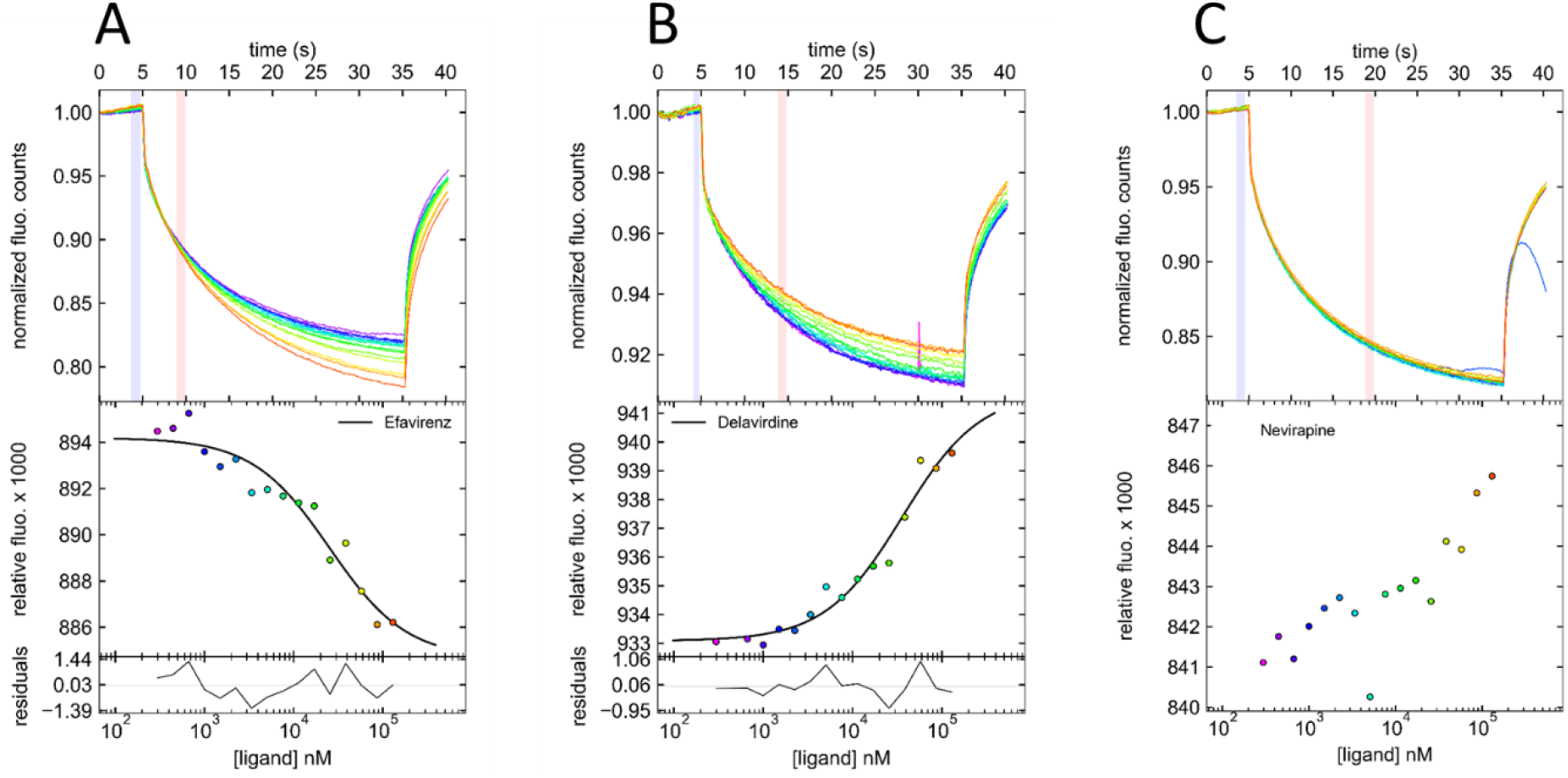
Microscale thermophoresis (MST) demonstrates a direct interaction between HIV-1 RT inhibitors and RED fluorescent-tagged TNF-α. For analysis, the change in thermophoresis is expressed as the change in the normalized fluorescence (Δ*F*_norm_), which is defined as *F*_hot_/*F*_cold_ (*F*-values correspond to average fluorescence values between defined areas marked by the red and blue cursors, respectively). Titration of the non-fluorescent ligand results in a gradual change in thermophoresis, which is plotted as Δ*F*_norm_ to yield a binding curve, which can be fitted to derive binding constants. **(A**) Experimental MST traces of efavirenz (*K*_D_ = 24 ± 8 μM); (**B**) Experimental MST traces of delavirdine (*K*_D_ = 39 ± 9 μM); (**C**) Experimental MST traces of nevirapine. Only the best MST traces (highest signal to noise ratio) are shown here. Values for all experiments conducted according to different experimental protocols are listed in **S2 Table**.

MST traces in presence of efavirenz and delavirdine showed distinct states (from bound to unbound), indicating a direct interaction of these two inhibitors with TNF-α (**Fig 3A, B**). Dissociation constants (*K*_D_) could be derived for the two corresponding complexes and estimated to 24 ± 8 μM (Efavirenz) and 39 ± 9 μM (Delavirdine), respectively (**Fig 3A, B**). The measured dissociation constants for the two HIV-1 RT inhibitors are in the same range of magnitude than that of UCB-6876 (A7M compound, K_D_= 22 μM; **Fig 2**), one of the three TNF-α inhibitors [22] used as a reference for this study.

Contrarily to our prediction, no thermophoresis signal could be detected in presence of nevirapine (**Fig 3C**) indicating no binding of this inhibitor to TNF-α, at least in our experimental settings. The herein observations were insensitive to experimental protocols (buffer composition, solubilizing agents, incubation time, MST power; **S2 Table**).

We next investigated the consequence of efavirenz and delavirdine binding to TNF-α, by looking at the binding of the cytokine to its cellular receptor (TNFR1) in presence of the two HIV-RT inhibitors. Indeed, both inhibitors were able to inhibit the binding of radiolabeled TNF-α to its receptor, albeit at much higher concentration (30-35% inhibition at 100 μM; **S1 Fig**) than the dissociation constants observed by MST experiments. Of course, none of the HIV-1 RT inhibitors has been optimized for binding to TNF-α and is directly repurposable for treating TNF-α −dependent autoimmune disorders. However, the recently discovered UCB6876 TNF-α inhibitor (A7M compound, **Fig 2**) [22], a fragment hit with a ligand efficiency similar to that of efavirenz (UCB6876: 0.34 kcal/mol per non-hydrogen atom; efavirenz: 0.32 kcal/mol per non-hydrogen atom) could be easily optimized to yield the 9 nM inhibitor UCB-5307 (A6Y compound, **Fig 2**) by just occupying a side pocket with a pyridyl ring [22]. Structure-guided efavirenz optimization for TNF-α binding is therefore possible by appropriate trimming of unnecessary cyclopropylethynyl substituent and occupation of the above potency subpocket.

### The similarity between TNF-α trimer and HIV-1 reverse transcriptase binding sites is not obvious

To demonstrate whether the herein disclosed similarity between the human TNF-α trimer and the HIV-1 RT non-nucleoside binding sites is obvious, we performed the same set of pairwise binding site comparisons, as that previously reported for ProCare (**Fig 1**), with state-of-the-art methods [10] developed either in-house (FuzCav [30], Shaper [21] and SiteAlign [31]) or by third parties (G-LoSA [32], KRIPO [15] and ProBiS [33]). The binding site perception, comparison algorithm and scoring function is specific to each method. Some methods (FuzCav, SiteAlign) consider entire cavities while some others utilize either fragment-bound subpockets (KRIPO, Shaper) or local protein descriptors (G-LoSA). To make the comparison consistent, the same set of atomic coordinates were compared, a binding site being defined by the protein PDB identifier, the ligand HET code and list of amino acids lining the cavity. The only exception was for the KRIPO method, which used all the chains available in the PDB entry. For each method, the distribution (**Fig 4**) and percentage (**Table 2**) of pairwise comparisons scored above the developer’s recommended similarity threshold were reported. Strikingly, only the G-LoSA method relying on a graph-based local alignment of cavity-lining amino acids, managed to find some similarity between the two sets of binding sites, however with reduced success rate (35.2%) when compared to the ProCare algorithm (76.6 % success rate; **Table 2**).

**Fig 4.**
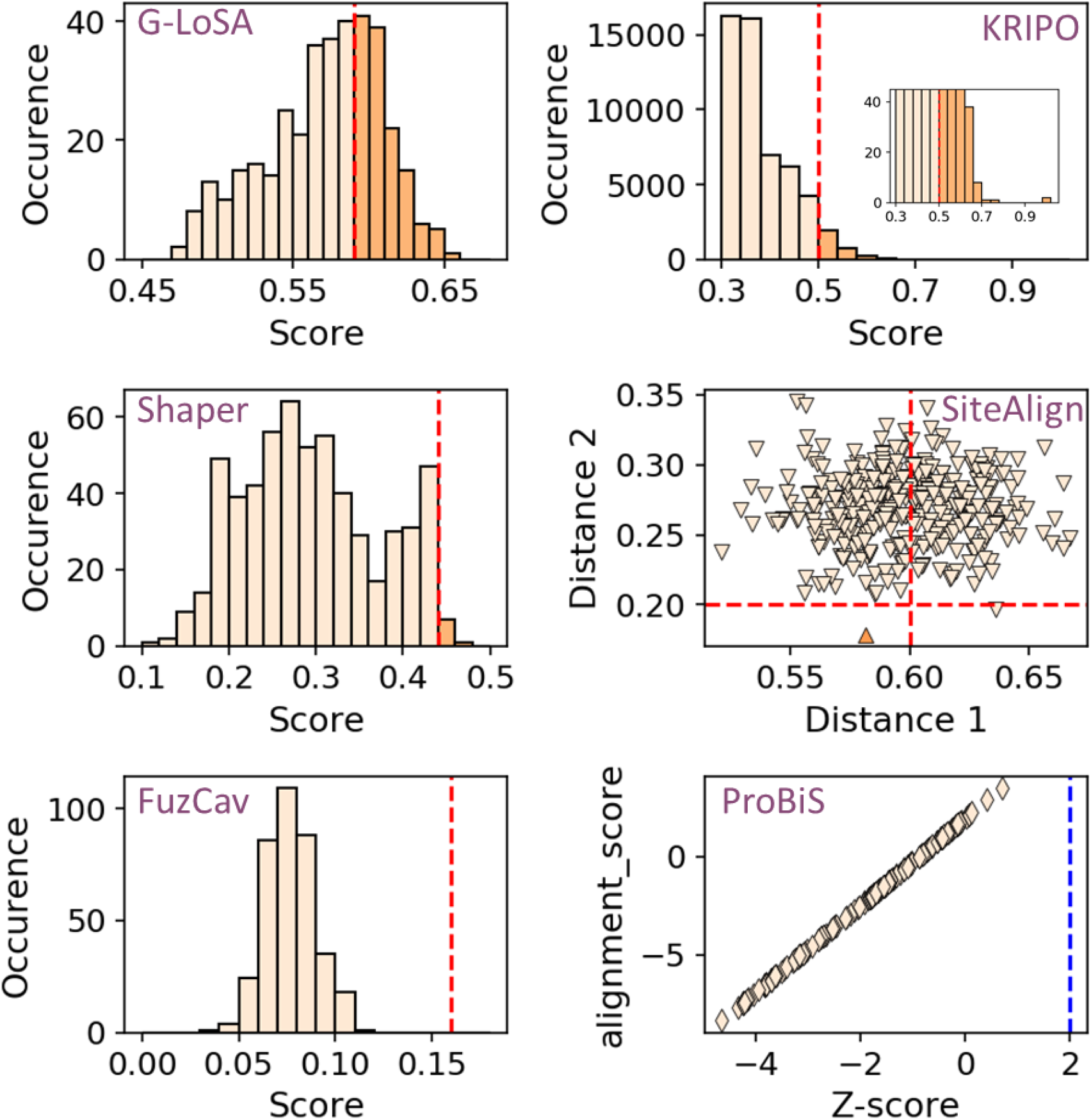
Score distribution of pairwise comparisons between binding sites of TNF-α trimer and HIV-1 reverse transcriptase. Binding sites in in asymmetric structures of TNF-α trimer (n=3) were compared to binding sites of non-nucleoside inhibitors in HIV-1 reverse transcriptase (sc-PDB set, n=122). Pairs with similarity measures scored above each method-specific threshold (red dashed line) were considered similar. For SiteAlign comparisons, pairs are considered similar in case the two distances (distance 1, distance2) are below the recommended cut-off. For ProBiS, the threshold above which an alignment is considered significant is marked by the blue dashed line.

**Table 2.** Comparison of three TNF-α and 122 HIV-1 RT non-nucleoside binding sites by state-of-the-art cavity comparison methods.

| Method | Score threshold<sup>a</sup> | Metric | Success rate<sup>b</sup> |
| --- | --- | --- | --- |
| G-LoSA | 0.59 | GA-score | 35.2 |
| KRIPO | 0.50 | Modified Tanimoto coefficient | 5.8 |
| Shaper | 0.44 | ColorRefTversky | 1.4 |
| SiteAlign | 0.6, 0.2 | d1 and d2 distances<sup>c</sup> | 0.3 |
| FuzCav | 0.16 | Tanimoto coefficient | 0 |
| ProBiS | 2 | Z-score<sup>d</sup> | 0 |
| ProCare | 0.47 | ProCare score | 76.6 |

<sup>a</sup> Developer's recommended similarity/distance threshold for estimating two binding sites similar
<sup>b</sup> Percentage of pairwise comparisons scored above the threshold.
<sup>c</sup> For SiteAlign comparisons, pairs are considered similar when the two distances (d1, d2) are below an acceptable cut-off.[31]
<sup>d</sup> The Z-score indicates the statistical relevance of ProBiS binding site alignments.

The herein reported binding of some HIV-1 RT non-nucleoside inhibitors to human TNF-α remains unobvious to many binding site comparison algorithms. Would this unexpected feature be better captured by remote ligand similarities? To investigate this question, we compared 2D and 3D descriptors of the corresponding inhibitors (**Fig 5**). Neither comparing 2D fingerprints nor 3D shapes would have confidently suggested possible binding of HIV-1 RT inhibitors to TNF-α trimer (**Fig 5**) since none of the considered ligand pairs exhibit a pairwise similarity above an acceptable threshold (Morgan2 circular fingerprint: 0.30 [34]; 166 public MACCS keys: 0.65 [34], TanimotoCombo ROCS 3D similarity: 1.5 [35–36]). We should precise here that 3D similarities were inferred from PDB protein-bound ligand X-ray structures and that alternative conformations might be selected by the two targets, although the very rigid efavirenz does indeed bind to the two proteins of interest albeit with different affinities. Extending 2D fingerprint comparisons to additional 2,361 HIV-1 RT inhibitors from the ChEMBL database [37], did not change our conclusion since only 0.71% and 0.09% of the corresponding pairs were found similar using Morgan2 and 166 public MACCS keys, respectively (data not shown).

**Fig 5.**
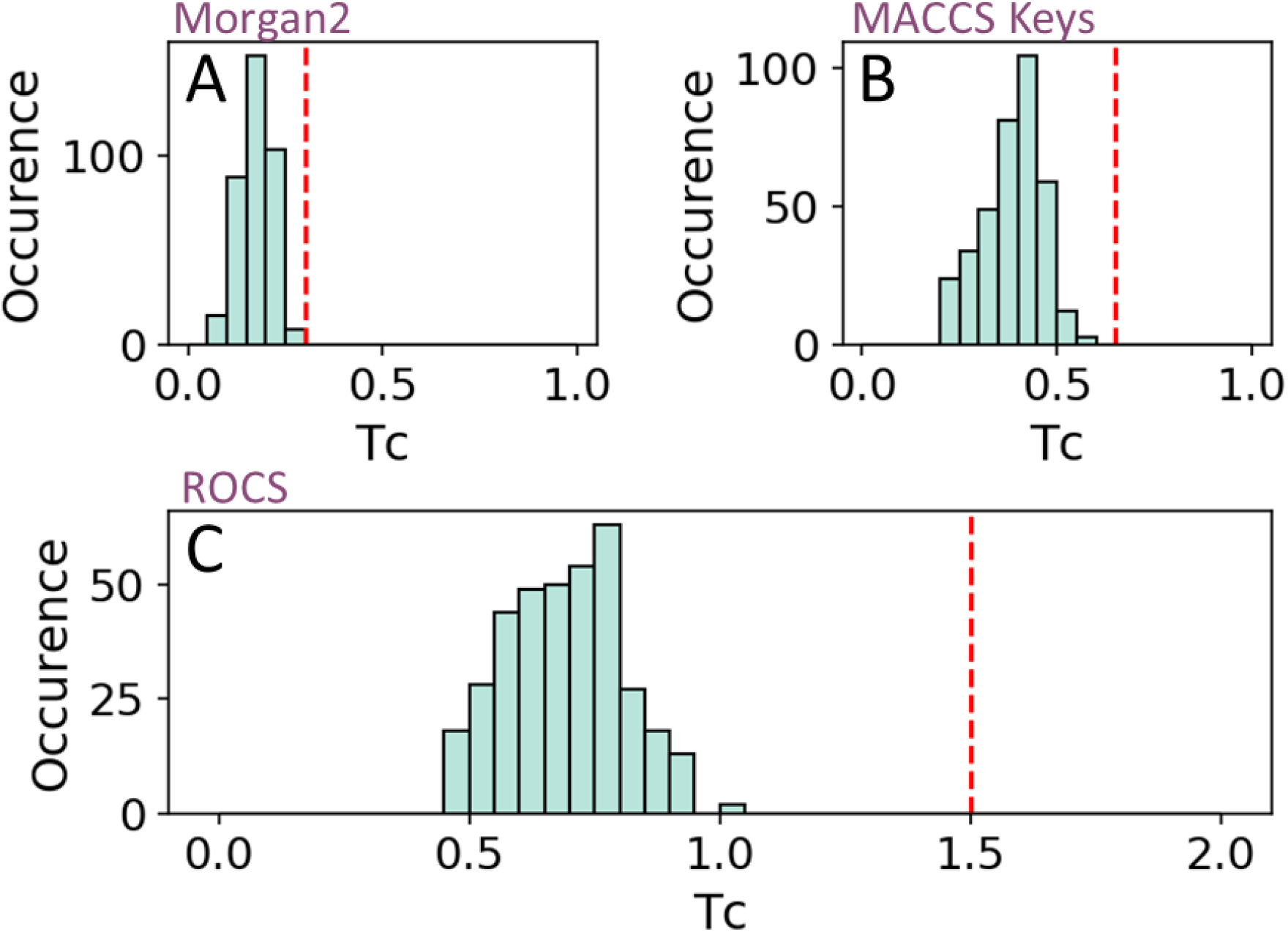
Pairwise similarity between inhibitors of TNF-α trimer and non-nucleoside inhibitors of HIV-1 reverse transcriptase. Recently described TNF-α trimer inhibitors (n=3) were compared to non-nucleoside inhibitors of HIV-1 RT (sc-PDB set, n=122). Pairs with similarity measures scored above each descriptor-specific threshold (red dashed line) were considered similar. (**A**) 2D similarity estimated by a Tanimoto metric using Morgan2 circular fingerprint, (**B**) 2D similarity estimated by a Tanimoto metric using 166 MACCS public keys. (**C**) 3D shape comparison (ROCS) estimated by the TanimotoCombo metric.

## CONCLUSION

Herein, we describe a systematic comparison of fragment-bound subpockets from *a priori* unrelated targets (TNF-α, HIV-1 RT) but predicted to share local similarities according to our recently-developed ProCare point cloud registration method. The computational prediction was verified by microscale thermophoresis experiments evidencing the micromolar binding of some but not all HIV-1 RT non-nucleoside inhibitors to human soluble TNF-α. Interestingly, the ProCare prediction could not be revealed by other state-of-the-art cavity or ligand similarity search methods. Point cloud registration, a computational method frequently used for digital image processing in robotics and medical imaging, enables the detection of subtle and local protein similarities thanks to a powerful description of subpocket microenvironments. The ProCare method appears as a promising idea generator for drug repurposing and fragment-based ligand design since it is able to pick starting ligands at a proteomic scale. Interestingly, the methods is not necessarily influenced by existing ligand or cavity neighborhoods.

## MATERIALS AND METHODS

### TNF-α dataset

The recently described asymmetric structures of the human TNF-α trimer bound to different inhibitors were retrieved from the RCSB Protein Data Bank (PDB) homepage (https://www.rcsb.org) [38] using the following identifiers: 6OOY, 6OP0, 6OOZ[22]. The PDB structures were protonated with Protoss [39] v.4.0, then split into protein, ligands and water molecules and finally converted into mol2 format with Sybyl-X v.2.1.1 (Certara USA, Inc., Princeton, NJ 08540). The binding sites (‘SITE’) were defined as any protein residue with at least one heavy atom closer than 6.5 Å from any ligand heavy atom and saved in mol2 and pdb formats. The ligands were converted into sdf format with OpenEye Python toolkits (OpenEye Toolkits 2020.2.0, OpenEye Scientific Software, Santa Fe) v.2020.0.4. ‘All’ VolSite [21] cavity points were computed with IChem v.5.2.9 [40] using default parameters.

### HIV-1 reverse transcriptase scPDB dataset

Starting from the PDB structure 1VRT as a reference, a search was performed in the RCSB PDB (https://www.rcsb.org) [38] to retrieve all structures with strict matching (“Structure Similarity” query in the PDB). After visual check, 122 entries already available in the sc-PDB repository (http://bioinfo-pharma.u-strasbg.fr/scPDB) [8] and for which the ligand is a non-nucleoside inhibitor were kept. The remaining PDB structures were protonated with Protoss [39] v4.0. The list of the PDB identifiers and Uniprot accession numbers are reported **S3 Table**. According to the sc-PDB preparation rules, the binding sites (‘SITE’) were defined as described above. Protein, ligand and binding site ‘SITE’ structures were directly retrieved in mol2 file format from the sc-PDB archive. The corresponding 122 ligands were 3D-fragmented with the IChem v.5.2.9 [40] fragmentation utility [25] and the complementary VolSite [21] cavity points, computed at 4 Å around each fragment were finally saved.

### HIV-1 reverse transcriptase ChEMBL dataset

Bioassay information were retrieved from the ChEMBL [37] dataset (Release 28; https://www.ebi.ac.uk/chembl) by querying the general keyword ‘reverse transcriptase’ and retaining ChEMBL target identifiers (CHEMBL247, CHEMBL4296301, CHEMBL2366516) corresponding to HIV-1 RT. Ligands with a measured sub-micromolar half-maximal inhibitory concentration (IC_50_) against the HIV1-RT single target were defined here as inhibitors (**S4 Table**). The corresponding SMILES strings were retrieved and further processed with RDKit (Open-source cheminformatics; http://www.rdkit.org) v.2019.03.4.0 to remove redundancy.

### Binding sites comparisons

#### ProCare

ProCare [17] v.0.1.1 pairwise comparison were performed on cavities computed with the VolSite module [21] in IChem v5.2.9 [40]. Entire cavities (“cavity_all” output) were calculated for TNF-α structures whereas only cavity points closer than 4.0 Å from any fragmented ligand heavy atom (“cavity_4” output) were considered for HIV-1 RT binding sites. VolSite cavity points were directly used for point cloud registration and determination of colored fast point feature histograms (c-FPFH) as previously described [17]. Finally, the respective set of c-FPFH descriptors for the two cavities were aligned and compared to each other using a RANSAC algorithm [19, 20] with default parameters [17]. Alignments results were scored with the default ProCare scoring function [17] which evaluates with a Tversky metric the proportion of aligned points of the same pharmacophoric features. In agreement with our previous study, a similarity ProCare score of 0.47 (p-value of 0.05) was used as threshold to distinguish similar from dissimilar binding sites.

#### FuzCav

FuzCav [30], an alignment-free method, was used to compare the binding site ‘SITE’ (mol2 format) entries of TNF-α dataset to the binding sites of HIV-1 RT sc-PDB dataset. Each binding site was tagged to compute a 4,833 bit-string that count all possible pharmacophoric triplets based on the atomic coordinates of Cα atoms lining the binding cavity. The pairwise comparisons of the fingerprints were evaluated with the default similarity score, with a threshold set at a value of 0.16 to distinguish similar from dissimilar binding sites.

#### G-LoSA

G-LoSA [32] v.2.2 is an alignment tool that was used with the binding sites ‘SITE’ pdb files. G-LoSA computes a set of inter-structural Cα pair distances to derive a graph, which will later be subjected to maximum clique search. The default G-LoSA score (GA-score) was used to evaluate the alignments. A threshold value of 0.59, recommended by the authors [32] and corresponding to a p-value of 0.05, was used to distinguish similar from dissimilar binding sites.

#### KRIPO

PDB ligands structural information were downloaded from Ligand Expo (http://ligand-expo.rcsb.org/) and prepared according to the KRIPO procedure (https://github.com/3D-e-Chem/kripo). Then KRIPO [15] v.1.0.1 was used with the list of prepared PDB structures for the pharmacophore fuzzy fingerprints calculations, using default parameters (fragmentation procedure activated). The pairwise similarities of the fingerprints were estimated with kripodb (v.3.0.0) using the modified Tanimoto coefficient as similarity metric. A threshold value of 0.50 was used to distinguish similar from dissimilar binding sites.

#### ProBiS

In a first place, the surface information (srf files) was computed for each prepared PDB structures with the default parameters referenced in the manual (3.0 Å to the ligand). ProBiS [33] requires a list of ligand HET code and residue number for each PDB entries. That list was provided to ensure that the ligands/sites considered are the same as in the binding site datasets used for other methods. Then, the alignment and comparison of the srf files were executed with default parameters, except for the Z-score that was set to a high negative value (−9999) as suggested by the authors to enforce the output of all results. Similarity between two binding sites was evaluated by the alignment score and Z-score. A threshold Z-score value of 2.0 was used to distinguish significant from irrelevant binding site alignments.

#### SiteAlign

For each entry, the list of natural amino acids in the ‘SITE’ mol2 files were provided as input. SiteAlign [31] v.4.0 describes a binding site by a polyhedron of 80 discretized triangles annotated with eight possible pharmacophoric features projected from cavity-lining C-α atoms. This results in a fingerprint of 640 integers. The pairwise comparisons imply aligning the corresponding polyhedron and computing the d1 and d2 distances of the fingerprints. The distance thresholds of d1=0.6 and d2=0.2 were applied respectively, to discriminate similar from dissimilar binding sites.

#### Shaper

Shaper [21] v.1.0 uses the same input files (VolSite cavities in mol2 file format) as ProCare. Shaper is an alignment method based on the OpenEye ShapeTK toolkit (OpenEye Toolkits 2020.2.0, OpenEye Scientific Software, Santa Fe) to maximize the overlap of shape and pharmacophoric features of the two compared cavities, thanks to a smooth Gaussian function. The alignments were realized with default settings and scored with a Tversky metric putting more weight on the reference cavity (RefTve). A threshold RefTve value of 0.44 (p-value = 0.005) was used to distinguish similar from dissimilar binding sites.

#### Ligand 2D similarity

Morgan fingerprints on the one hand, and 166 public MACSS keys on the other hand were computed on the sc-PDB ligands (sdf format) and ChEMBL ligands (SMILES strings) with RDKit python package v.2019.03.4.0 using default parameters (radius = 2 Å for the Morgan fingerprints). The Tanimoto coefficients of the pairwise TNF-α ligands/HIV-1 RT ligands fingerprints comparison were reported. Cut-off values of 0.30 (Morgan fingerprints) and 0.65 (MACCS keys) were used to discriminate chemically similar from dissimilar ligands.

#### Ligand 3D similarity

sc-PDB HIV-1 RT inhibitors were compared to TNF-α inhibitors with OpenEye ROCS v.3.2.0.4 and scored by decreasing Tanimoto similarity metric accounting for both shape and chemical features overlap (TanimotoCombo). A TanimotoCombo cut-off value of 1.5 was used to discriminate chemically similar from dissimilar ligands.

##### Chemicals and biologicals

Nevirapine (catalog #S1742), efavirenz (catalog #S4685) and delavirdine mesylate (catalog #S6452) were purchased from Selleck Chemicals (https://www.selleckchem.com/). Soluble human TNF-α (catalog # Z01001) was purchased from GenScript (http://www.genscript.com).

##### Binding of HIV-1 RT inhibitors to human TNF-α (Microscale thermophoresis)

Human TNF-α was labeled using the RED-NHS 2nd generation labeling kit (NanoTemper Technologies GmbH) using a protein concentration of 10 μM and a molar dye-to protein ratio ~ 3:1. A label/protein ratio of 0.4 was determined using photometry at 650 and 280 nm. Compounds efavirenz, delavirdine and nevirapine were initially dissolved in DMSO to afford stock solutions of 10 mM. These were then diluted to initial concentrations of 260 μM into 20 mM K phosphate pH 7.4, 150 mM NaCl ensuring a final concentration of DMSO of 2.6 %. These compounds were serially diluted 2:1 in buffer 20 mM K phosphate pH 7.4, 150 mM NaCl, 2.6 % DMSO producing ligand concentrations ranging from 260 μM to 594 nM (16 titration points). For MST measurements, each ligand dilution was mixed with 1 volume of fluorescently-labelled TNFα at 680 nM in 20 mM K phosphate pH 7.4, 150 mM NaCl, 0.02% Tween-20, which leads to a final concentration of TNFα of 340 nM and final ligand concentrations at half of the ranges above. The final buffer is 20 mM K phosphate pH 7.4, 150 mM NaCl, 0.01% Tween-20 and 1.3 % DMSO. After a 15-min incubation at room temperature in the dark, followed by centrifugation at 13,000 g for 3 min, each solution was filled into Monolith NT Premium capillaries (NanoTemper Technologies GmbH). Thermophoresis was measured at 25°C with 40% LED power and 20, 40% and 80% MST power using a Monolith NT.115 (NanoTemper Technologies GmbH). Data were analysed in the NT Analysis software version 1.5.41 (NanoTemper Technologies GmbH).

## SUPPORTING INFORMATION

**S1 Fig. Inhibition of 0.1 nM [125I]-TNF-α binding to human TNF receptor type 1 (TNFR1) in U-937 cells (Brockhaus et al. Proc. Natl. Acad. Sci. USA, 87, 3127-3131, 1990), by three HIV-1 reverse transcriptase inhibitors (Eurofins Discovery assay #76).** Results are mean ± sem of two experiments.

(TIF)

**S1 Table. sc-PDB Pockets found similar to the inner cavity of human TNF-α trimer(PDB ID 6OOY).**

(XLSX)

**S2 Table. Dissociation constant (KD) of three HIV-1 RT inhibitor binding to human soluble TNF-α, according to MST experimental conditions.**

(XLSX)

**S3 Table. List of PDB entries describing non-nucleoside inhibitors bound to HIV-1 reverse transcriptase.**

(XLSX)

**S4 Table. List of CHEMBL entries describing non-nucleoside inhibitors.**

(XLSX)

**S1 Video. Alignment of nevirapine subpocket (PDB ID: 1VRT, HET code: NVP, color: purple) to the TNF-α pocket (PDB ID: 6OOY, HET code: A7M, color: white).**

(MP4)

## ACKNOWLEDGMENTS

The authors are thankful to the Doctoral School of Chemical Sciences (EDSC, University of Strasbourg) for a doctoral fellowship to M.E. The Calculation Center of the IN2P3 (CNRS, Villeurbanne, France) is acknowledged for the allocation of computing time and excellent support. We sincerely thank Prof. M. Rarey (University of Hamburg, Germany) for providing an executable version of Protoss, OpenEye Scientific Software for the generous allocation of an academic license, and Dr. Catherine Birck (IGBMC, Illkirch) for the microscale thermophoresis experiments.

## AUTHOR CONTRIBUTIONS

Conceived and design experiments: ME, DR. Performed the experiments: ME, Analyzed the data: ME, DR. Wrote the paper: ME, DR.

